# Genome dynamics across the evolutionary transition to endosymbiosis

**DOI:** 10.1101/2023.05.02.539033

**Authors:** Stefanos Siozios, Pol Nadal Jimenez, Tal Azagi, Hein Sprong, Crystal L Frost, Steven R Parratt, Graeme Taylor, Laura Brettell, Kwee Chin Liew, Larry Croft, Kayla C King, Michael A Brockhurst, Václav Hypša, Eva Novakova, Alistair C Darby, Gregory DD Hurst

## Abstract

Endosymbiosis – where a microbe lives and replicates within a host – is an important contributor to organismal function that has accelerated evolutionary innovations and catalysed the evolution of complex life. The evolutionary processes associated with transitions to endosymbiosis, however, are poorly understood. Here, we use comparative genomics of the genus *Arsenophonus* to reveal the complex processes that occur on evolution of an endosymbiotic lifestyle. We compared the genomes of 38 strains spanning diverse lifestyles from environmentally acquired infections to obligate inter-dependent endosymbionts. We observed recent endosymbionts had larger genome sizes than closely related environmentally acquired strains, consistent with evolutionary innovation and rapid gain of new function. Increased genome size was a consequence of prophage and plasmid acquisition including a cargo of type III effectors, and concomitant loss of CRISPR-Cas genome defence systems enabling mobile genetic element expansion. Persistent endosymbiosis was also associated with loss of type VI secretion, likely reflecting reduced microbe-microbe competition. Thereafter, the transition to stable endosymbiosis and vertical inheritance was associated with the expected relaxation of purifying selection, pseudogenisation of genes and reduction of metabolism, leading to genome reduction. However, reduced %GC that is typically considered a progressive linear process was observed only in obligate interdependent endosymbionts. We argue that a combination of the need for rapid horizontal gene transfer-mediated evolutionary innovation together with reduced phage predation in endosymbiotic niches drives loss of genome defence systems and rapid genome expansion upon adoption of endosymbiosis. These remodelling processes precede the reductive evolution traditionally associated with adaptation to endosymbiosis.

## Results and Discussion

Animals live in a microbial world. Their interactions with microbes range from the antagonistic with pathogenic symbionts, through to the mutualistic with beneficial symbionts, with evolutionary transitions occurring commonly between these states [1, 2]. Transitions in symbiotic interactions can further select for evolution of key symbiotic traits, such as vertical transmission and eventual integration of symbionts into host anatomy and physiology. These transitions correspondingly alter selection pressures. For example, vertical transmission relaxes selection for traits necessary for external survival whilst also correlating microbe transmission with host fitness and thus favouring beneficial function(s). Concurrently, population bottlenecks associated with vertical transmission limit within-host symbiont diversity, selecting for lower virulence [3]. Further, these bottlenecks intensify genetic drift [4], reducing the efficiency of purifying selection for function. These processes collectively drive genome degradation through pseudogenization, genome reduction and lowered %GC [5].

Endosymbiosis is the state in which the symbiont lives within the body or cells of the host organism. Evolutionary transitions from environmental to endosymbiotic lifestyles are common across the tree of microbial diversity, and have occurred with microeukaryotic, fungal, plant and animal hosts [2]. However, our understanding of the evolutionary processes associated with transitions to endosymbiosis is limited. In particular, there have been few studies of the initial transition to host association and vertical transmission inherent to becoming an endosymbiont. Fewer chart the entire transition from environmentally acquired associations through to obligate vertically inherited endosymbiont via a facultative endosymbiotic stage.

To gain a precise view of the tempo and mode of evolution during the transition to persistent endosymbiosis, we leveraged the wide diversity of host-associated lifestyles in the gammaproteobacterial genus *Arsenophonus*. Within this clade, there are environmentally acquired extracellular pathogens, extracellular endosymbionts with mixed modes of transmission, intracellular facultative endosymbionts with vertical transmission, and obligate vertically transmitted endosymbionts where the partners are co-dependent [6-10]. Importantly, extracellular pathogens and endosymbionts with mixed modes of transmission are very closely related. This genus thus provides a unique opportunity to investigate the genomic adaptations enabling the emergence of an endosymbiotic host association, building from work in other clades with more coarse-grained comparisons [11, 12].

To gain a high-resolution view of the evolutionary transition to endosymbiosis in an insect-associated bacterial endosymbiont, we first completed closed genomes for seven new strains: three *Arsenophonus nasoniae* from *Nasonia vitripennis*, one from each of the parasitic wasps *Pachycrepoideus vindemmiae* and *Ixodiphagus hookeri*, one from the blue butterfly *Polyommatus bellargus*, and a strain of *Arsenophonus apicola* isolated from Australian *Apis mellifera*. We also completed a draft genome for *Ca*. A. triatominarum. In addition, novel draft genomes for a further 17 *Arsenophonus* strains were assembled from Sequence Read Archive (SRA) deposits from a variety of insect genome sequencing projects. Genome assembly data, alongside sample details and current understanding of transmission mode and nature of symbiosis, are given in Supplementary Table 1.

We then estimated the phylogenetic affiliation of the strains through core genome analysis (Figure 1), maximizing the common gene set for phylogenetic inference by excluding the strains with highly reduced genomes. This analysis revealed three main clades according to lifestyle; the *nasoniae* clade where characterized members have mixed modes of transmission or have recently become facultative vertically transmitted symbionts; the *apicola* clade where characterized strains are infectiously transmitted, and the *triatominarum* clade where characterized members are vertically transmitted. When the obligate co-dependent symbionts are included in the phylogeny (Supplementary Figure S1), they do not form a monophyletic clade as was previously shown [13], indicating independent evolutionary transitions to obligate host dependence. Culturable representatives are common in the *apicola*/*nasoniae* clades, likely reflecting a current or recent requirement to live outside the endosymbiotic environment. Members of the bee genus *Bombus* have members from both *apicola* and *nasoniae* clades, indicating recurrent colonization by the endosymbiont of this host group.

**Figure 1.**
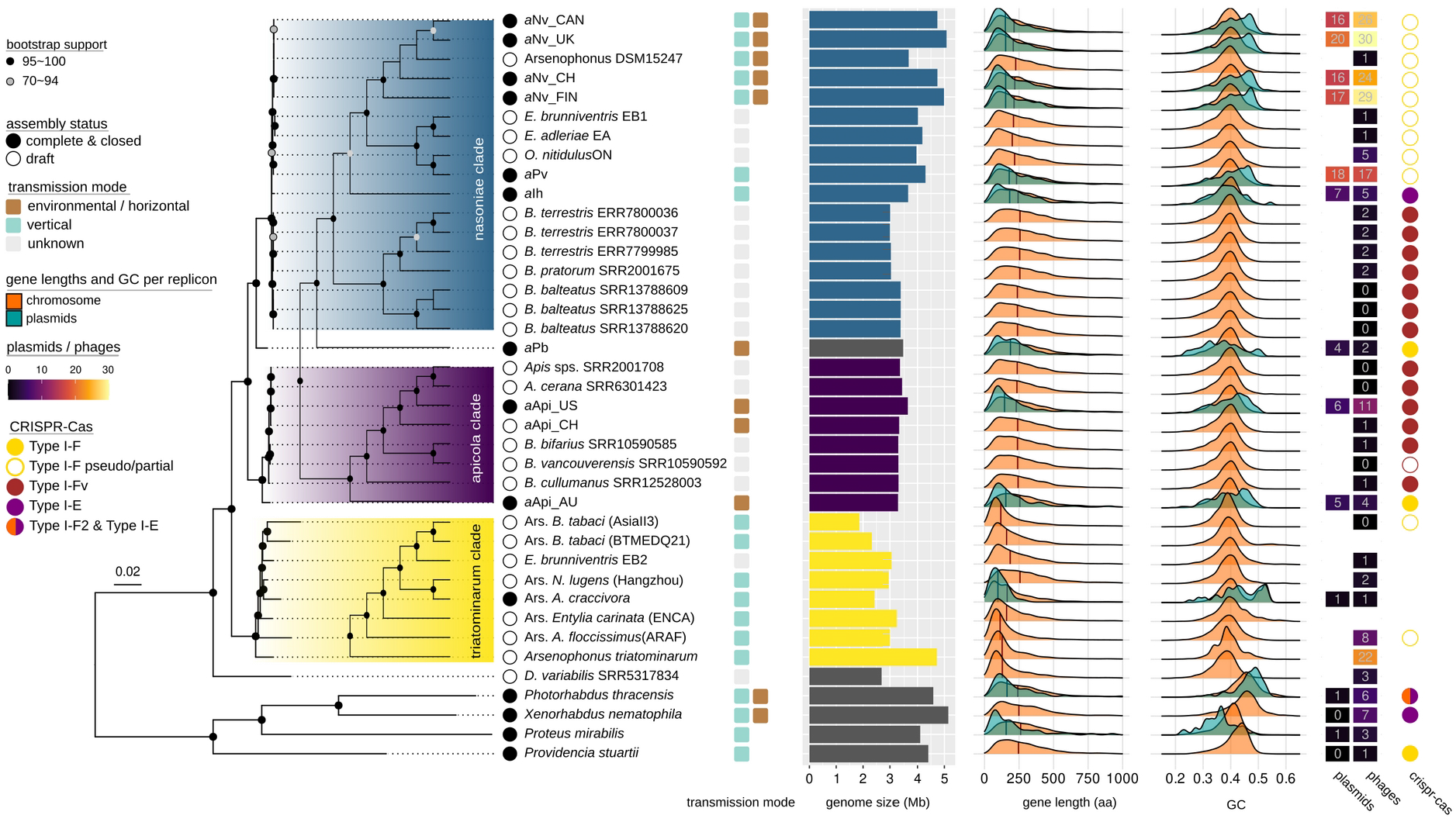
Core genome phylogeny and genome features of the Arsenophonus clade. The phylogenetic relationships between the Arsenophonus strains were inferred using maximum likelihood on the concatenated set of 230 single copy core protein sequences in IQ-TREE v2.1.4 under the JTTDCMut+F+R3 model. Only bootstrap support values >= 70 are shown. Inset cladograms are used to improve tree readability. The nasoniae, apicola and triatominarum clades are highlighted in blue, magenta and yellow respectively, while complete genomes are indicated by black circles in front of the tip labels. The transmission mode of each strain is indicated with the coloured squares (green: vertical, brown: horizontal and grey: currently unknown). The horizontal bar-plot shows the genome size in MB. The ridgeline plots show the distribution of CDS length (in amino acids) and GC content fraction for genes of chromosomal (orange) or extra-chromosomal (green) origin. The vertical lines in the gene length ridgeplot represent median values. The heatmap shows the number of plasmids and intact phages, while the coloured circles indicate the type and intactness of the CRISPR-Cas system (filled circle: intact, non-filled circle: pseudogenised, no circle: not present). A Bayesian phylogenetic analysis including the four obligate Arsenophonus genomes (Arsenophonus of Lipoptena fortisetosa, Arsenophonus of Aleurodicus dispersus, Arsenophonus of Ceratovacuna japonica and Arsenophonus of Melophagus ovinus) and Ca. Riesia is shown in the Supplementary Figure S1.

*Arsenophonus* strains varied in genome size from 663,125 to 5,080,918 bp. It is generally considered that symbiont genome size is an inverse function of time evolving as endosymbiont under vertical transmission, with strains with long history of vertical transmission having smaller genomes. As expected, the smallest genomes in the clade were indeed from interdependent obligate endosymbionts and members of the triatominarum clade where all members are vertically transmitted and require a host for replication (Figure 2A). Unexpectedly, the five strains with the largest closed genomes were not those with environmental transmission, but related strains with either mixed modes of transmission or vertical transmission. This pattern reflected larger genome size in the nasoniae clade (where characterized members have protracted host association and vertical transmission and genome size ranges from 3.65-5.1MB) compared to the apicola clade (where characterized members are environmentally acquired pathogens and genome size is 3.27-3.63 Mb) (Figure 2A). The increase in genome size in early-stage endosymbionts is driven largely by an increase in mobile genetic elements, notably prophage and plasmid content, in strains which have recently entered endosymbiosis and lost environmental transmission (Figure 1).

**Figure 2.**
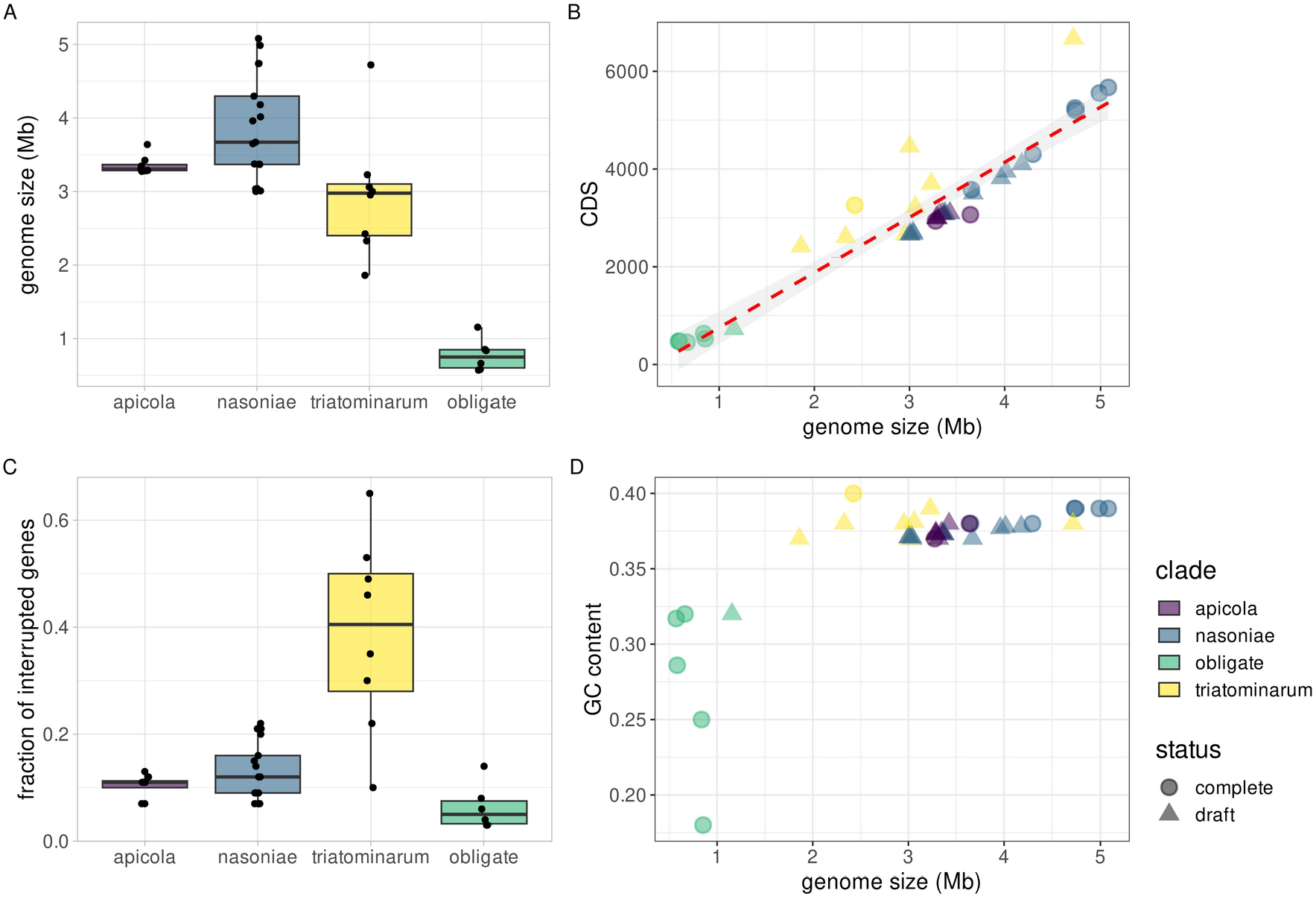
Genomic characteristics of the Arsenophonus/Riesia clades. **A**) Genome size distribution across Arsenophonus clades, **B**) Association between genome size and the number of coding sequences (CDS). The red dashed line in panel B represents a fitted linear trend line with confidence intervals shown as grey shading. **C**) the fraction of interrupted genes across Arsenophonus clade. This was estimated by calculating the fraction of proteins with length <80% of the length of their top hit in the Swiss-Prot database. **D**) Association between genome size and the fraction of GC content. Although the number of predicted protein-coding genes shows, as expected, a linear relationship with the genome size across the Arsenophonus/Riesia clades, this association does not hold for GC content contrary to the classical observations between free-living and symbiotic microbes. Box plots: center line, median; box limits, 25th and 75th percentiles; whiskers, ±1.5x interquartile range; data points are shown with the black dots.

Acquisition of mobile genetic elements, like prophages and plasmids in bacteria, is often accompanied with horizontal gene transfer of accessory genes that are important in microbial virulence and adaptation [14, 15]. Consistent with this pattern, we observed a gain in Type III secretion (T3SS) associated effectors in nasoniae group strains that have recently become endosymbionts, compared to environmentally acquired strains (Figure 3). These data support previous work in *Sodalis* arguing symbionts repurpose the T3SS on transition from pathogenesis to enable persistent intracellular host association [16]. Further to this hypothesis, our data indicate that the process may in some cases involve acquisition of new effectors. Contrastingly, T3SS are either absent or heavily pseudogenized in the triatominarum clade where vertical transmission is well established. The loss of T3SS systems in highly derived vertically inherited endosymbiont genomes suggests that these traits become redundant or costly once the host and endosymbiont have become tightly coevolved.

**Figure 3.**
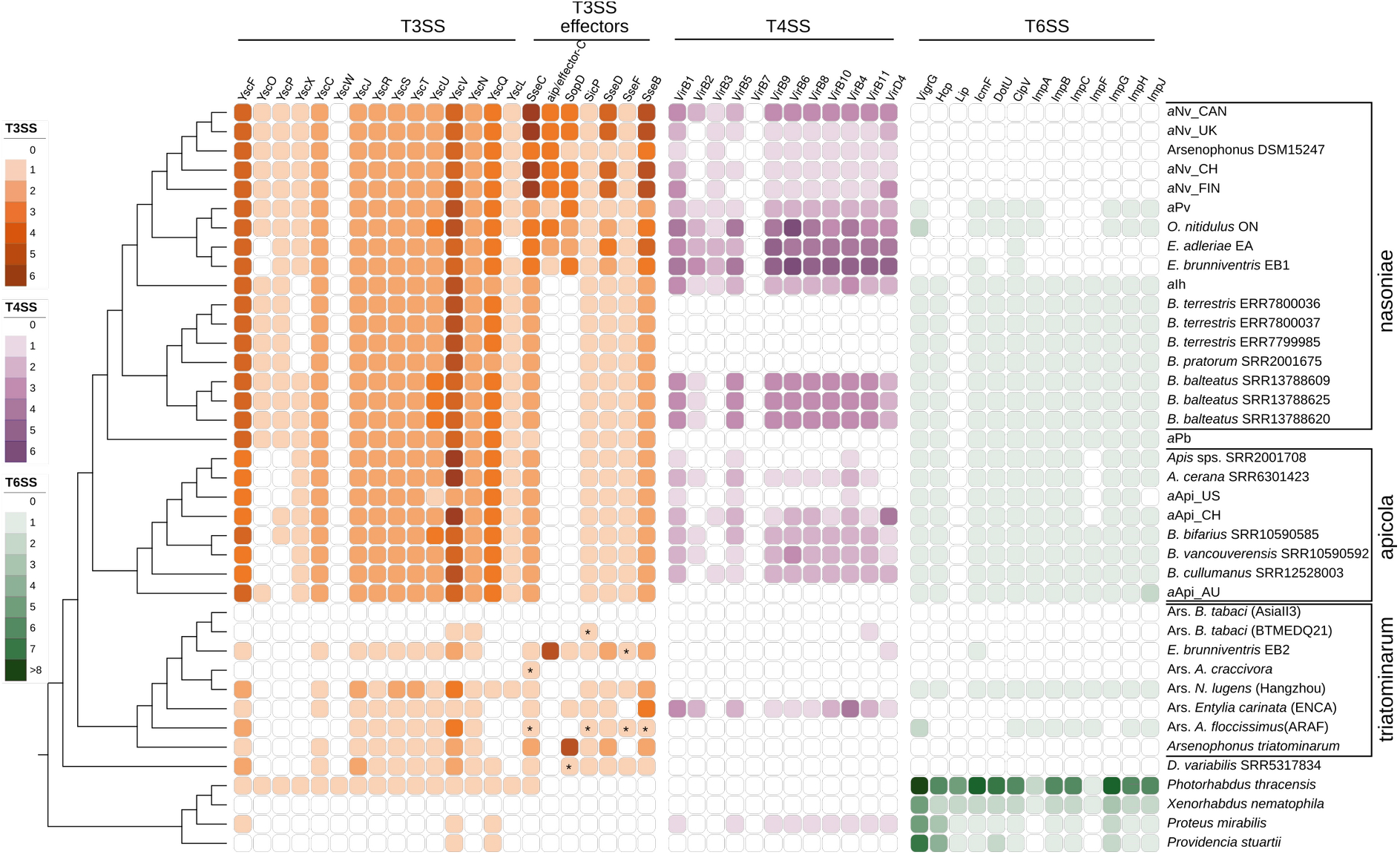
Comparative analysis of secretion systems across the Arsenophonus clades. Core components of the Type III (orange), type IV (magenta) and type VI (green) systems as predicted by BlastKOALA are shown. The absence of genes is indicated by empty squares. Identified type III effectors are also shown. The relationship of the Arsenophonus strains is shown with the cladogram based on the core genome phylogeny (Figure 1). The asterisks indicate potential pseudogenes. Not shown in the Figure: Secretion systems are absent in the obligate Arsenophonus lineages (Arsenophonus of Lipoptena fortisetosa, Arsenophonus of Aleurodicus dispersus, Arsenophonus of Ceratovacuna japonica and Arsenophonus of Melophagus ovinus) and Ca. Riesia.

We next investigated potential drivers of prophage and plasmid accumulation. Past analysis has indicated that genome defence systems, such as CRISPR-Cas systems which protect the genome from mobile DNA, are less commonly found in symbiotic microbes than in free-living ones [17]. We observed that the identity, distribution and completeness of CRISPR-Cas genome defence systems varies extensively among *Arsenophonus* genomes (Figure 4). We identified three types of CRISPR-Cas systems (types I-F, I-Fv and I-E) with variable gene content across all strains, suggesting dynamic turnover of these important genome defence systems (Figure 4A). This turnover is likely mediated by horizontal gene transfer and recombination (Supplementary Figure S2), consistent with variable mobile genetic element-mediated selection for genome defence across niches. Variable mobile element exposure is further supported by the distinct lack of spacer matches between different *Arsenophonus* clades. This result suggests that different clades have been exposed to distinct mobile genetic element communities (Figure 4B).

**Figure 4.**
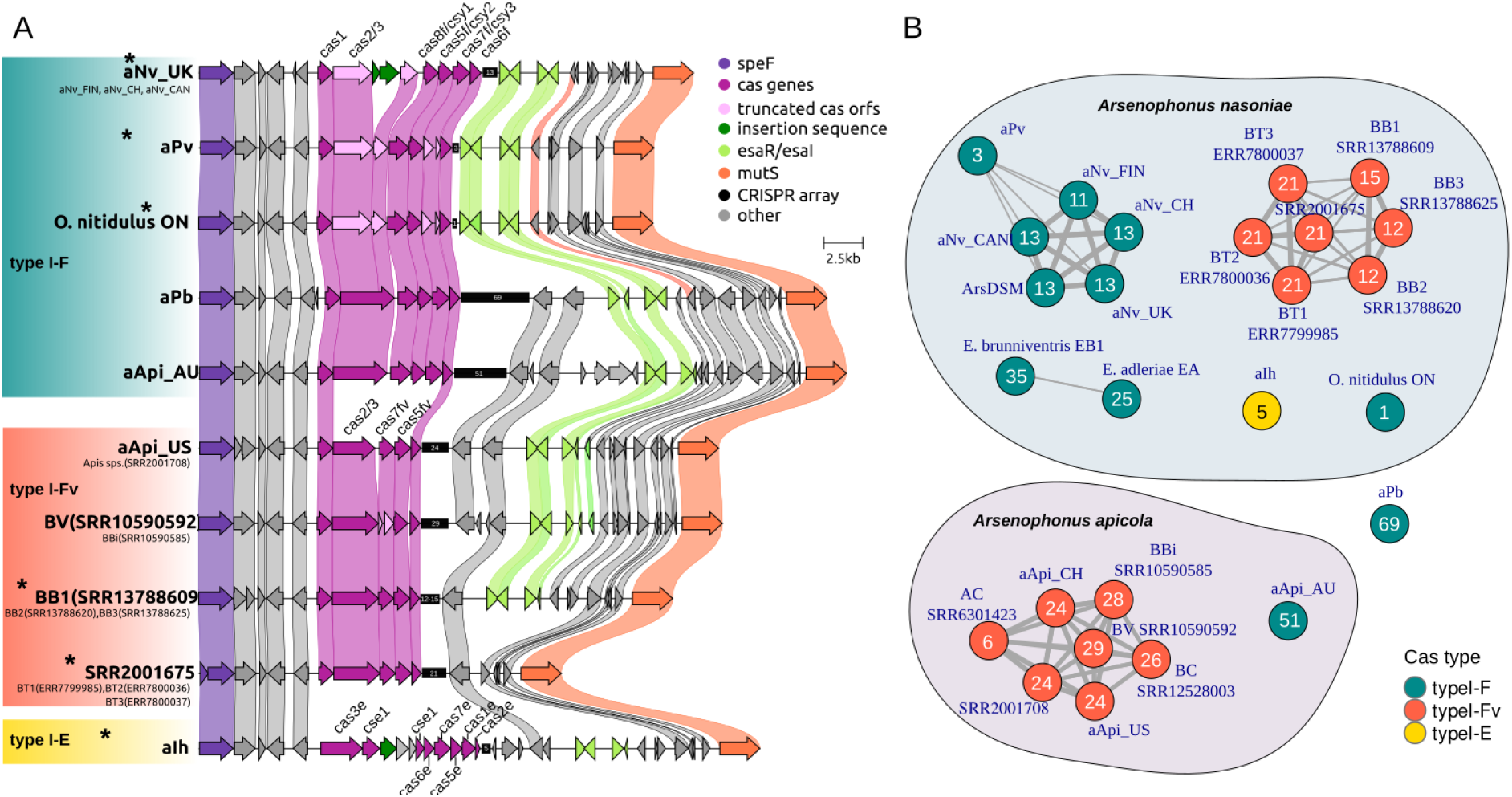
The CRISPR-Cas phage defence system in Arsenophonus and early signs of pseudogenisation within the nasoniae clade. **A**) Gene order and genomic context of the Arsenophonus CRISPR-Cas systems plotted with clinker software. Groups of homologous genes are connected by coloured ribbons. Arsenophonus strains belonging to the nasoniae clade are highlighted with an asterisk. **B**) Networks showing the relatedness of Arsenophonus genomes based on shared CRISPR spacers. Nodes correspond to genomes and the edges are scaled based on the spacer repertoire relatedness (see details in Methods section). Absence of edge correspond to no shared spacers between the genomes. Nodes are coloured according to the Cas type identified in each genome. The numbers in the nodes represent the number of spacers identified by CRISPRCasFinder tool.

Intact CRISPR-Cas systems were predominantly found in environmentally acquired strains. Notably, closely related vertically inherited strains contained recently pseudogenized CRISPR-Cas systems (evidenced by intact and shorter CRISPR arrays but insertions within the Cas systems), while obligate intracellular symbionts carried either highly degraded fragmentary CRISPR-Cas systems or none (Figure 1 and Figure 4A). CRISPR-Cas is known to have metabolic and autoreactive costs [18]. Theoretical and experimental data suggest that rapid loss of CRISPR-Cas function is under selection as a means to maintain horizontally transferred genetic elements that are beneficial for microbial adaptation [19, 20]. Thus, the extensive pseudogenization and loss of CRISPR-Cas systems outside of environmentally acquired strains likely reflects purifying selection against maintaining this genome defence systems within host environments. Selection against CRISPR-Cas within host-associated environments could reflect the selection to retain beneficial mobile genetic elements whilst avoiding autoimmunity or may be due to lower rates of phage attack within hosts. Aside CRISPR-based defences, other predicted anti-phage systems are also more diverse in the environmentally acquired strains compared to strains with mixed modes of transmission. These are fragmentary in strains exhibiting vertical transmission (Supplementary Figure S3 and Supplementary Table S2).

Within intracellular and other endosymbiotic host-associated niches, bacteria are likely to encounter fewer competitors. To test if this weakened selection to retain anticompetitor weapons, we examined the presence and integrity of type VI secretion systems (T6SS), a contact-dependent system for killing competitor microbes [21]. Whereas T6SS were present in the environmentally acquired strains, these were incomplete in nasoniae clade strains with either mixed modes or vertical transmission and absent or fragmentary in all but one member of the triatominarum clade. It is in this last clade where all members show vertical transmission (Figure 3). T6SS are multiprotein complexes and thus likely to be costly to produce and use ([22] but see [23]). Their loss in endosymbionts supports a model where reduced competition within the endosymbiotic niche reducing the benefits of maintaining T6SS.

Classically, the transition to endosymbiosis has been associated with reduced metabolic capability of endosymbionts. The stable nutritional environment within host cells reduces the need for metabolic plasticity. Contrastingly, a recent study of Chlamydiae concluded that the transition to intracellular endosymbiosis was associated with increased predicted metabolic capability of the symbionts [24]). In the *Arsenophonus* clade, predicted metabolic functions are reduced compared to free living ancestors. However, metabolic capabilities are not markedly distinct between vertically transmitted endosymbionts and environmentally-acquired strains. The completeness of metabolic pathways broadly reflects the abilities of strains to grow in *in vitro* cell-free culture [9]. One notable exception is beta oxidation of fatty acids, which was only intact in environmentally acquired strains; this result was sustained when draft genomes for the triatominarum group were additionally examined (Supplementary Figure S4). Aspects of cofactor synthesis, amino acid synthesis and carbohydrate metabolism were degraded in strains that live intracellularly, and loss of function in these pathways was most pronounced for obligate co-dependent endosymbiont strains. Together, these patterns suggest that the transition to intracellular endosymbiosis, rather than endosymbiosis *per se*, is associated with evolved losses of metabolic functions. Whether this degradation is a function of longer evolutionary time under relaxed selection or is actively selected for by the intracellular niche is unclear.

We further examined gene loss processes over the life course of symbiosis. As expected, the number of predicted CDS was an approximately linear function of genome size (Figure 2B). The fraction of pseudogenized genes was low in both environmentally acquired strains and in interdependent obligate symbionts, at intermediate levels in the nasoniae group containing vertically transmitted strains that retain infectious transfer and culturability, and at highest levels in unculturable facultative endosymbiont strains (Figure 2C). This pattern was only apparent in non-core genes (genes not present in all strains) (Supplementary figure S5). These data suggest a process of pseudogenization that accelerates on entering into endosymbiosis, as systems for environmental survival become redundant in the host context. The number of pseudogenes then increases over time until purifying selection drives their loss and genomes become streamlined to core genes.

A key hypothesis in symbiont evolution is that vertical transmission leads to bottlenecks in symbiont population size. This process is expected to increase the importance of drift over selection and thus weaken the capacity of purifying selection to maintain function [25]. We analysed the pattern of molecular evolution of highly conserved single copy genes that are critical for microbial function (see methods for details). All three clades (nasoniae, apicola and triatominarum) showed evidence of relaxed selection compared to the autonomous free-living outgroup. Relaxed selection was most pronounced in the triatominarum clade comprising vertically transmitted intracellular symbionts with a long history of endosymbiosis. Relaxation of selection was also observed in the other two clades. Notably, it was somewhat more pronounced in the nasoniae clade where the strains have mixed modes of transmission or vertical transmission than the apicola clade where environmental transmission is common (Figure 5). These data corroborate our current thinking of evolution patterns in vertically transmitted symbiosis, which combines adaptive gene loss through redundancy with gene degradation through fixation of mildly deleterious alleles, the latter permitted by increased primacy of drift processes. Notably, this signature can be detected shortly following the evolution of vertical transmission.

**Figure 5.**
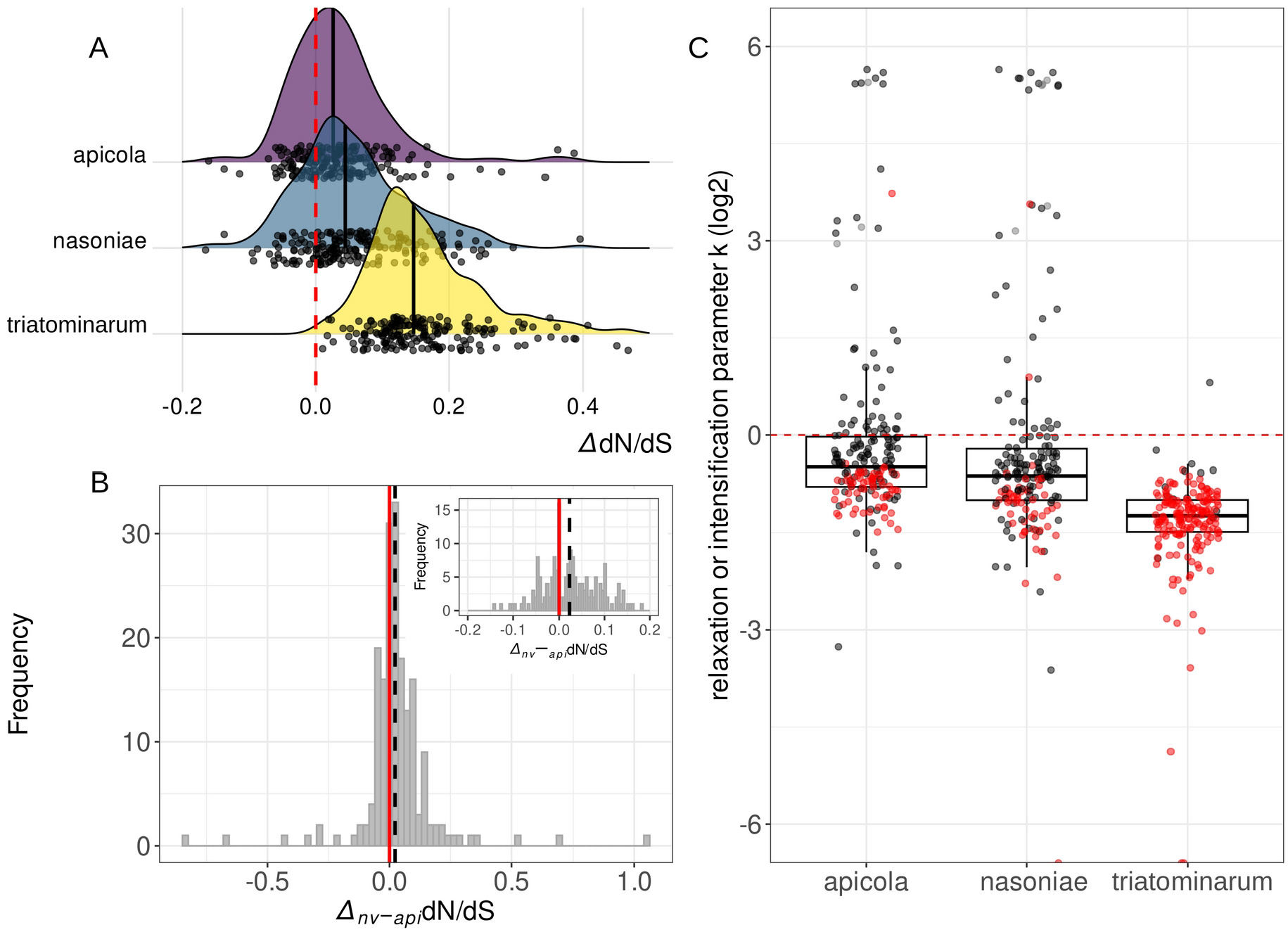
Evidence of relaxation of selection between Arsenophonus clades with contrasting modes of transmission. **A**) Distribution of gene-wise dN/dS ratios in the three main Arsenophonus clades as compared to the outgroup clade comprised of free-living species (Providencia stuartii, Proteus mirabilis, Morganella morganii and Moellerella wisconsensis) (dN/dS_test clade_ – dN/dS_outgroup_) for 188 highly conserved single-copy BUSCO marker genes. Individual values are shown as jitter points. Black solid lines represent the median of the distribution. In all clades the median is shifted to the right with the triatominarum clade (vertical transmission) showing the largest shift followed by the nasoniae clade (mixed mode of transmission) and last the apicola clade (environmental transmission) suggesting a gradual increase in the dN/dS ratios as we progress towards protract symbiosis. **B**) Distribution of gene-wise differences in dN/dS ratios between nasoniae and apicola clades for the same 188 highly conserved BUSCO marker genes. The black dotted vertical line represents the median of the distribution which is shifted to the right (median = 0.0228146, Wilcoxon signed rank test V = 11373, P = 4.018e-06) indicating that nasoniae clade has higher dN/dS ratios compared to apicola clade which is mostly characterised by environmental mode of transmission. A narrower range of the same data between values -0.2 and 0.2 is shown in the inset plot on the top right corner. **C**) Distribution of relaxation or intensification parameter k (log2) per gene as calculated by the RELAX method in HyPhy package (v2.3) compared to the outgroup clade. Values below zero indicate that selection strength has been relaxed while values above zero indicate an intensification of the selection strength. Genes with statistically significant k values (FDR; q<0.1) are shown as red jitter dots.

Finally, we examined how overall genomic features vary between strains. Unlike previous work [26], our data do not support the linear association between genome size and %GC content during the transition to endosymbiosis, at least in the genus *Arsenophonus* (Figure 2D). Reduced %GC is only markedly observed in the obligate co-dependent endosymbionts with highly reduced genomes. For the other strains, %GC content is relatively consistent at 37-40%. Analysis of coding sequences of non–obligate strains did not support a relationship between %GC and genome size (Null hypothesis of no association: F_1, 33_ = 1.141, p=0.29; see Supplementary Figure S6). Our data suggest therefore that reduced %GC may be restricted to obligate interdependent symbioses in this group. This may also reflect changes in DNA repair systems which are ablated in obligate strains (Supplementary figure S7), increased primacy of genetic drift associated with the pronounced bottlenecks that accompany obligate interdependent symbioses, or a combination of these factors.

In summary, our examination of genome evolution across symbiotic lifestyles in the genus *Arsenophonus* reveals a new model for the evolution of endosymbiosis. Becoming a persistent endosymbiont requires rapid evolutionary innovation fuelled by horizontal gene transfer, notably the gain of new functions for host manipulation. This rapid genome expansion is achieved through the acquisition of prophage and plasmid mobile genetic elements, which is enabled by the loss of genome defence systems, including CRISPR-Cas. An initial establishment of endosymbiosis phase is associated with enrichment for T3SS effector toxins but this is followed by their loss in strains that become vertically transmitted and more highly adapted to the host intracellular environment. Whilst our model is based on data from a single genus, the centrality of CRISPR defence loss reflects recent studies of *Mycoplasma* evolution following a host switch event [27], and the presence of intact or recently pseudogenized CRISPR in culturable, but not unculturable, aphid-associated *Serratia* [12]. In *Mycoplasma*, the authors argued the loss of CRISPR systems enabled adaptation to the novel host species. We argue that the evolutionary transition from free-living to endosymbiosis may commonly be associated with a more complex genome dynamics than previously reported, with genome expansion and remodelling preceding the processes of reductive evolution traditionally associated with endosymbiosis.

## Method details

### Isolate collection, culture, sequencing and assembly

Seven *Arsenophonus* isolates were isolated to pure culture and sequenced within this project: *A. nasoniae* isolates aNv_UK, aNv_CH and aNv_CAN derived from *Nasonia vitripennis* from the UK, Switzerland and Canada respectively, *A. nasoniae aPv* from *Pachycrepoideus vindemmiae* and the *Arsenophonus* strain *a*Pb from the butterfly *Polyommatus bellargus* [9], *Arsenophonus nasoniae* strain *a*Ih previously identified in the parasitoid wasp *Ixodiphagus hookeri* from a questing *Ixodes ricinus* tick in The Netherlands [28] and *Arsenophonus apicola* strain aApi_AU from Australian honey bees described in [29]. A further strain, *Ca*. A. triatominarum, was isolated into cell culture. Details of isolation, culture, sequencing and assembly are given in Supplementary methods.

### *Arsenophonus* genomes assembled from publicly available SRA deposits

We screened publicly available SRA datasets (Source: DNA, Platform: Illumina, Strategy: genome) originated mainly from the Apoidea superfamily (containing bees and bumblebees) as well as Parasitoida infraorder and Ixodida order for the presence of *Arsenophonus* reads. We did that by performing a “*Mash screen*” using Mash v2.3 [30, 31] to measure the containment of a local database of reference Arsenophonus genomes within the unassembled SRA read sets. SRA datasets with at least 80% containment were taken for downstream processing. An initial metagenomic assembly of the short reads from the identified SRA datasets were performed using MEGAHIT v1.2.8 [32] and the assembled contigs >= 1.5kb were binned based on their deferential tetranucleotide frequencies using MetaBAT2 v2.12.1 under default parameters [33]. The *Arsenophonus* bins were identified and completeness was assessed using CheckM v1.0.18 [34]. To identify *Arsenophonus* contigs potentially missed from the initial binning process the original contigs from the metagenomic assembly were screened using blastn (-task megablast) against a local database consisted of all available and complete *Arsenophonus* genomes using BLAST 2.12.0+ [35]. Contigs >1kb with significant matches (e-value < 1e-25) to *Arsenophonus* genomes were extracted and included in the metabat bins. The augmented bins were quality inspected and further refined in anvio v7 [36] by identifying and removing potential contaminant contigs based on atypical coverage and gene-level taxonomic classification. The original BioProject and SRA accessions from which the draft Arsenophonus was obtained are shown in Supplementary Table S3.

### Comparative analysis of the metabolic potential across the *Arsenophonus* clades

To avoid inconsistencies stemming from draft and incomplete genomes, only the metabolic potential of complete *Arsenophonus* genomes was estimated. To these we included for comparison the genomes of the closely related and obligate symbionts *Ca*. Riesia pediculicola and *Ca*. Riesia pediculischaeffi as well as the genomes of the four outgroup species used in the phylogenetic analysis. All genomes were annotated for functions and metabolic pathways on the basis of KEGG database using the anvi-run-kegg-kofams command in anvio v7. KEGG MODULE metabolism was finally estimated using the anvi-estimate-metabolism pipeline as described in [37]. A KEGG module was considered to be complete in a given genome when at least 75% of the steps involved were present.

### Phylogenomic analysis

For the maximum likelihood phylogenomic analysis the highly divergent genomes from the obligate Arsenophonus strains (Ca. *Arsenophonus lipoptenae, Arsenophonus* of *Aleurodicus dispersus, Ca*. Arsenophonus melophagi and *Arsenophonus* of *Ceratovacuna japonica*) including *Ca*. Riesia pediculola were excluded as their placement can be affected by strong compositional heterogeneity and long branch attraction. The phylogenetic relationship of the remaining 35 *Arsenophonus* genomes was estimated on the concatenated set of 230 single-copy core protein sequences representing highly conserved gammaproteobacterial BUSCO v4.1.4 markers [38]. These were identified through Orthofinder v2.3.11 [39]. Alignments of individual protein orthologs were performed using mafft program [40] as implemented in Orthofinder and screened for recombination based on the Pairwise Homoplasy Index (PHI) test using the Phipack package [41] revealing no significant evidence. The alignments were quality trimmed using ClipKIT alignment trimming tool v1.3.0 [42] under the *smart-gap* mode before concatenated into a super matrix using seqkit [43].

Best protein model (JTTDCMut+F+R3) was identified using ModelFinder [44] and a ML phylogenetic tree was reconstructed in IQ-TREE v1.6.12 [45]. The genomes from the related species *Proteus mirabilis* strain HI4320 (NC_010554), *Providencia stuartii* strain MRSN 2154 (NC_017731), *Photorhabdus thracensis* strain DSM 15199 (NZ_CP011104) and *Xenorhabdus nematophila* ATCC 19061 (NC_014228) were used as an outgroup. All genomes were pre-annotated using prokka v1.13 [46] for consistency.

To more precisely estimate the relationships between the *Arsenophonus* strains including the obligate and highly diverse lineages a separate Bayesian phylogenetic analysis was conducted based on the concatenated set of 77 manually curated single-copy core protein clusters using PhyloBayes-MPI v1.9 [47] and the CAT-Poisson model. Briefly, the alignments of all single copy protein clusters (101) were manually inspected to minimize alignments with high gap content before concatenation. Two independent chains were run in parallel for at least 30,000 cycles until convergence was observed (rel diff < 0.1 and minimum effective size > 300 for all trace file metrics) assessed by running bpcomp and tracecomp commands in PhyloBayes.

### Annotation of genomic features

The proportion of pseudogenised genes were estimated by calculating the fraction of interrupted proteins using the Ideel method against the UniProt/Swiss-Prot database (https://github.com/mw55309/ideel). Prophage regions were annotated using the PHAge Search Tool Enhanced Release (PHASTER) web server [48]. Annotation of CRISPR arrays and cas genes was performed with the CRISPRCasFinder program using the default parameters [49]. CRISPR spacer relatedness was calculated as the number of shared spacers by two genomes divided by the number of spacers in the smallest array [50]. Shared spacers were identified based on an all-vs-all blastn search allowing for at least 98% similarity and 97% coverage between individual spacers. Apart from CRISPR-Cas systems we screened for additional phage defense systems using DefenseFinder [51].

### Analysis of relaxation of selection strength between Arsenophonus clades

We search for signatures of relaxation of selection on a set of 188 highly conserved single-copy BUSCO orthologs across 39 *Arsenophonus* strains using the HyPhy (Hypothesis Testing using Phylogenies) software package version v2.5.8 [52]. The obligate and highly diverged *Arsenophonus* strains including *Ca*. Riesia were excluded from the analyses. Four, mostly environmental, Morganellaceae (Enterobacterales) species (*Proteus mirabilis, Providencia stuartii, Morganella morganii* and *Moellerella wisconsensis*) were used as reference/outgroup since *Photorhabdus thracensis* and *Xenorhabdus nematophila* that were used in the phylogenetic analyses above are having complex modes of transmission involving both vertical and horizontal transmission. To this end, single-copy protein clusters were identified as before using Orthofinder v2.3.11 and 188 clusters representing highly conserved gammaproteobacterial BUSCO v4.1.4 markers were selected for downstream analyses. Alignment of protein sequences was performed with mafft using the “-auto” option and back translated to nucleotide alignments using pal2nal [53]. A phylogenetic tree was estimated using the JTTDCMut+F+I+G4 model in IQ-TREE v1.6.12 on the concatenated data of the same set of 188 orthologues protein clusters. This tree was then used to identify genes with significant evidences of relaxation or intensification of selection using the RELAX hypothesis testing framework in HyPhy package [54]. Three sets of branches were selected as the “test branches” (nasoniae clade, apicola clade and the triatominarum clade, see Supplementary Figure S8) and compared to the reference/outgroup set of branches to estimate the selection intensity parameter k for each gene. A value of k > 1 indicates that selection on the test branches is intensified compared to the reference branches while a value < 1 indicates a relaxation of selection. Statistically significant values were assessed through a likelihood ratio test (LRT) followed by a false discovery rate (fdr) correction to account for multiple comparisons. Branch specific dN/dS ratios were estimated for each individual gene using the partitioned MG94xREV model which fits a single dN/dS value to each branch partition as implemented in RELAX method in HyPhy. The differences of dN/dS ratios between branches (ΔdN/dS) were statistically assessed using a Wilcoxon signed-rank test in R v4.2.2 [55].

### Data visualization

Tools used for data visualization: R v4.2.2 [55], ggplot2 [56], patchwork [57], igraph v1.4.0 [58]. A presence/absence heatmap for metabolic pathways was generated using the pheatmap v1.0.12 (https://rdrr.io/cran/pheatmap/) package in R v4.2.2. Phylogenetic trees were drawn and annotated using the ggtree package v4.2 [59]. The gene order comparison of CRISPR-Cas systems was visualized in clinker v0.0.24 [60].

## Data and code availability

All genomes and the raw reads generated in this study are deposited in GenBank database under the BioProject accession number PRJNA956975. The genome for *Ca*. Arsenophonus triatominarum can be found in GenBank under the BioProject accession number PRJNA311587. The code and source data for the various analyses in this study can be found on GitHub (https://github.com/SioStef/Arsenophonus-comparative-genomics).

