## Supplementary Figures for "Genome dynamics across the evolutionary transition to endosymbiosis"

**Supplementary Figure S1.** Bayesian inference of *Arsenophonus* phylogeny including the highly divergent genomes from the obligate *Arsenophonus* strains (*Arsenophonus* of *Lipoptena fortisetosa*, *Arsenophonus* of *Aleurodicus dispersus, Arsenophonus* of *Melophagus ovinus* and *Arsenophonus* of *Ceratovacuna japonica*) including *Ca*. Riesia.

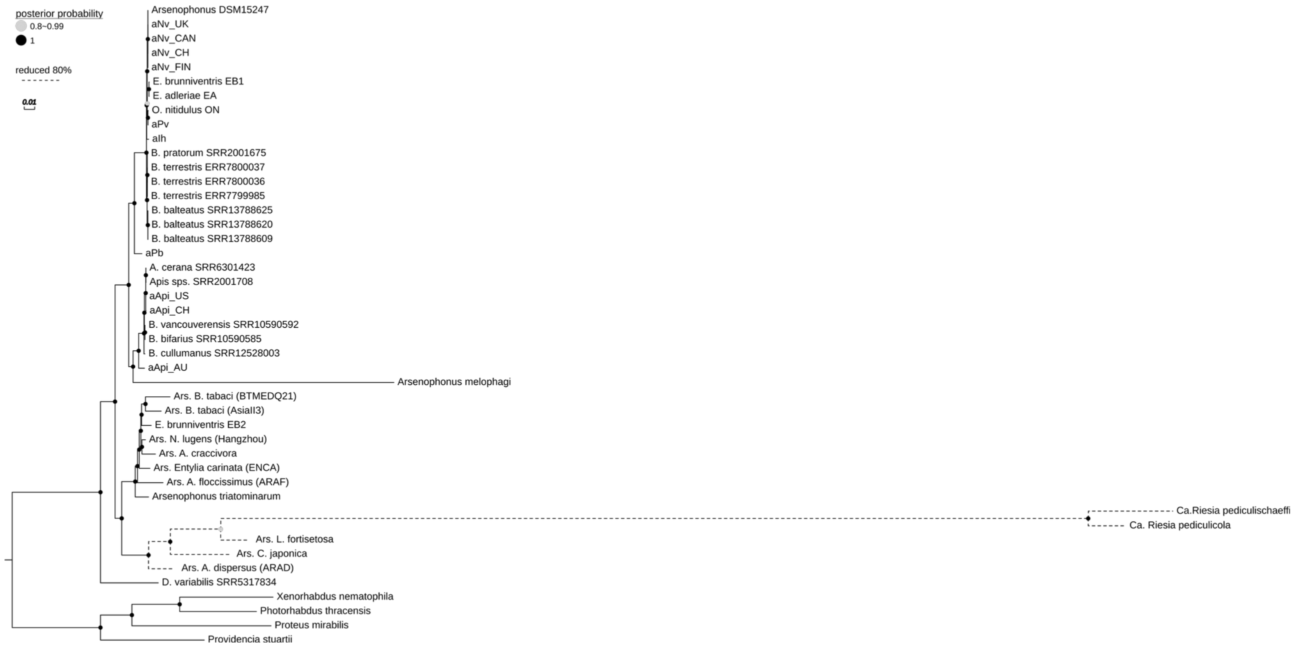

**Supplementary Figure S2.** Evidence of recombination within the CRISPR-Cas region. **A**) maximum likelihood phylogenies of two Cas proteins (Left: Cas1, Right: Cas6/Csy4). Sequences from the nasoniae and apicola clades are highlighted in blue and purple respectively. The incongruence between the two phylogenies suggest that recombination and horizontal gene transfer mediates the evolution of the *Arsenophonus* CRISPR-Cas system **B**) reticulated evolution of Cas1 proteins. The phylogenetic network was reconstructed from the protein alignment of Cas1 homologs using the SplitDecomposition method with SplitsTree version 4.19.0. Bootstrap values ( >95% ) based on 1000 replicates are indicated.

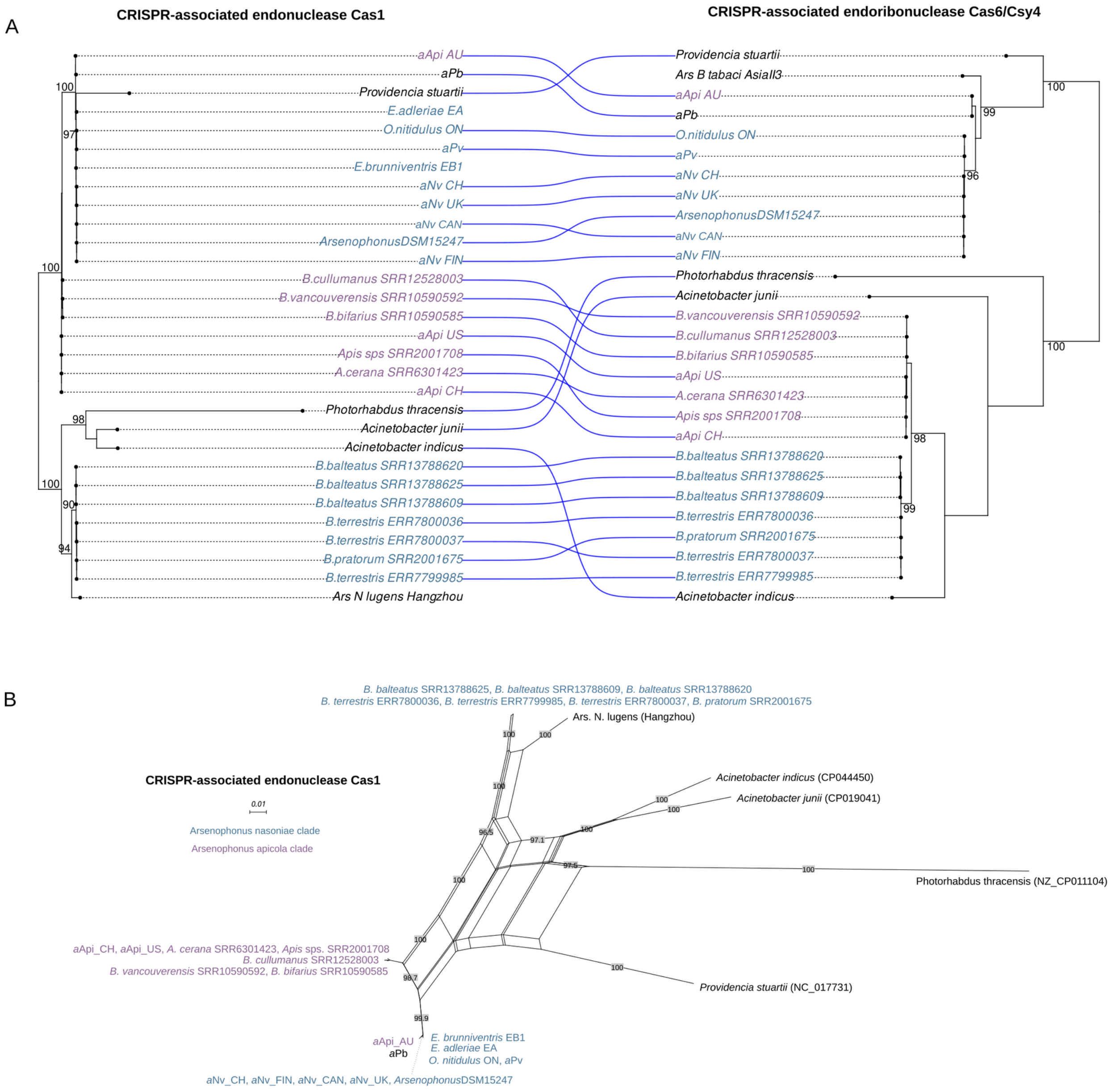

**Supplementary Figure S3.** Summary of predicted phage defence systems, other than CRISPR-Cas across the *Arsenophonus* clades. Obligate lineages are devoid of phage defence systems (not shown). The alternative phage defence systems were detected using DefenseFinder (Tesson et al. 2022).

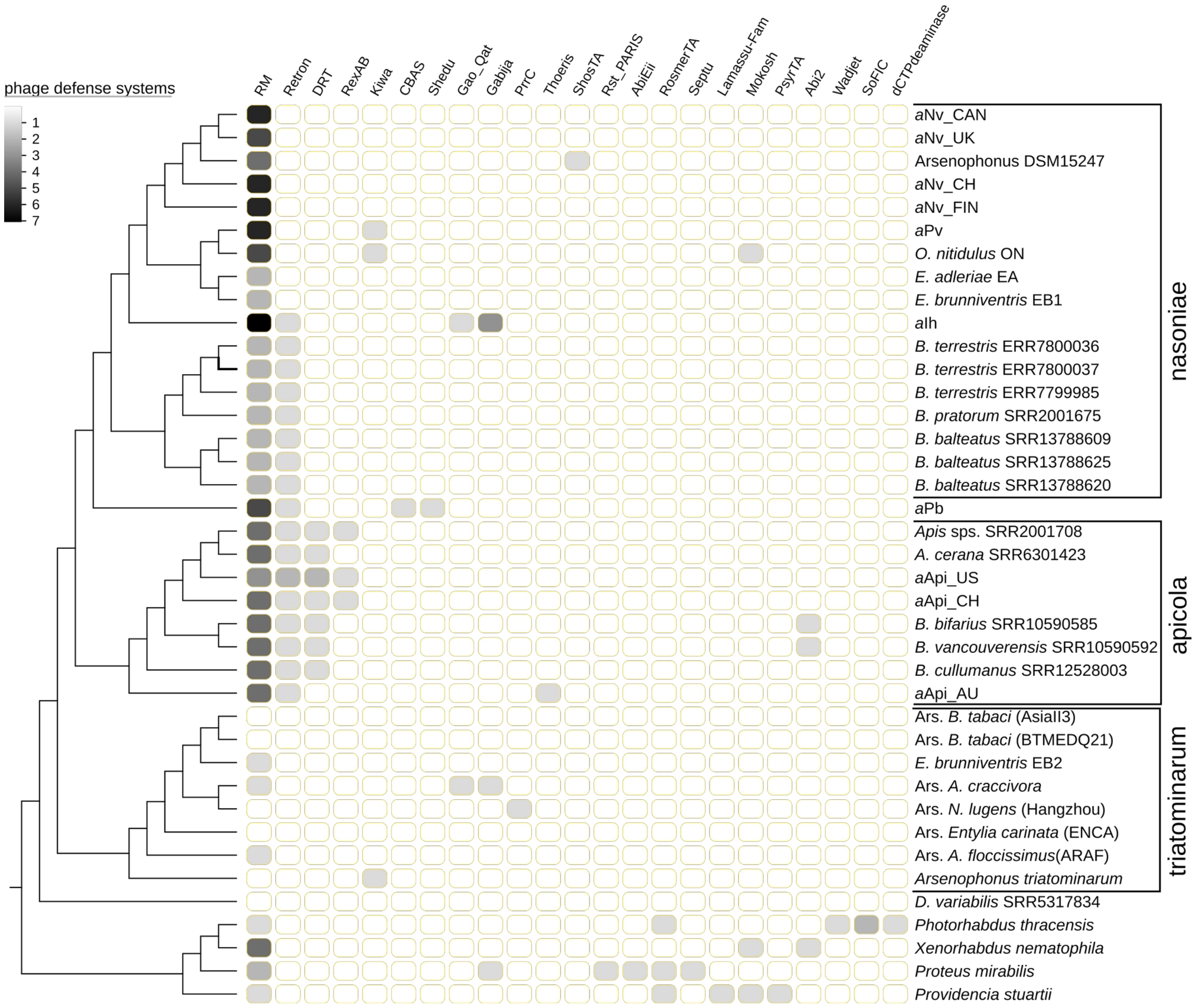

**Supplementary Figure S4** Comparison of the metabolic potential across *Arsenophonus*/*Ca*. Riesia lineages with complete genomes and selected outgroup species. KEGG MODULE metabolism was estimated in anvio version 7. A KEGG module was considered to be complete in a given genome when at least 75% of the steps involved were present.

A) Carbohydrate and energy metabolism

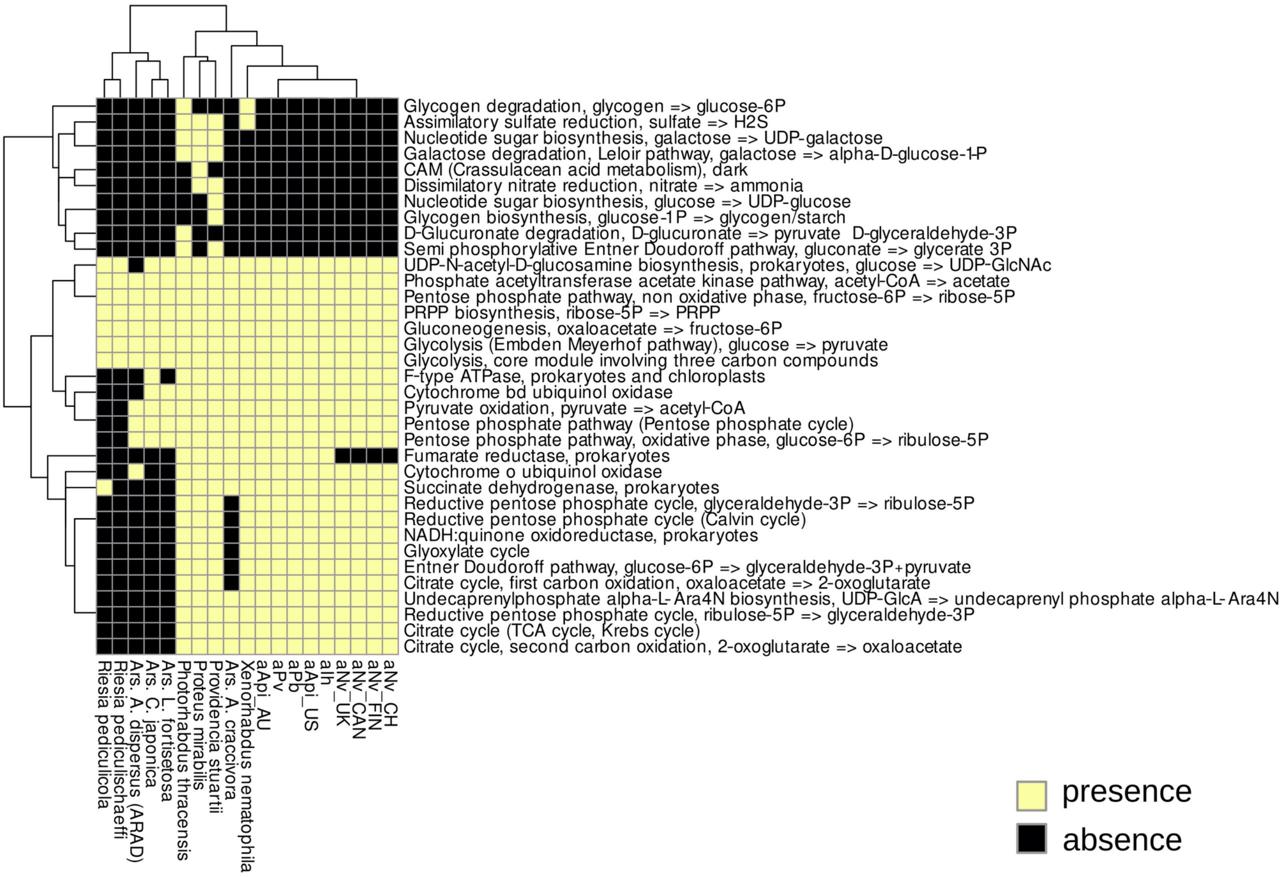


B) Lipid and glycan metabolism

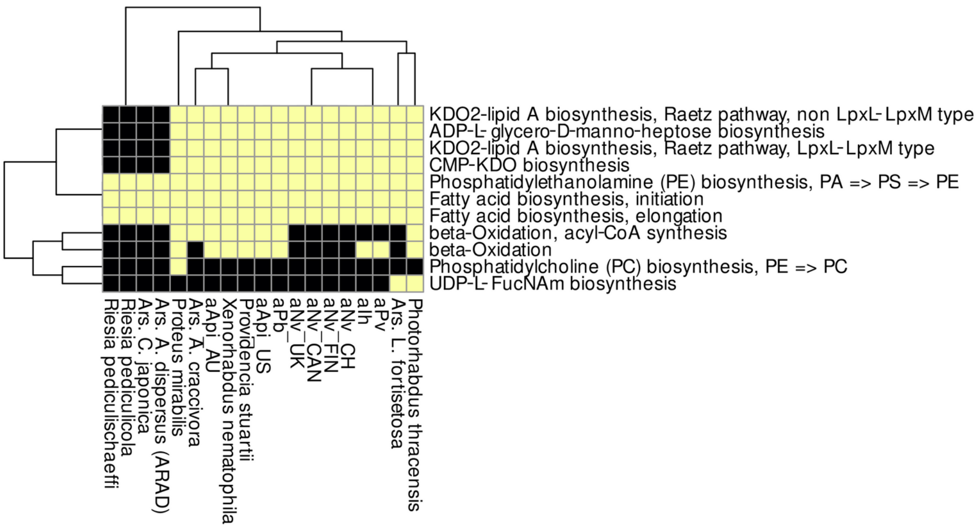

**Supplementary figure 4** …*(Continued from previous page)*

C) Amino acid metabolism

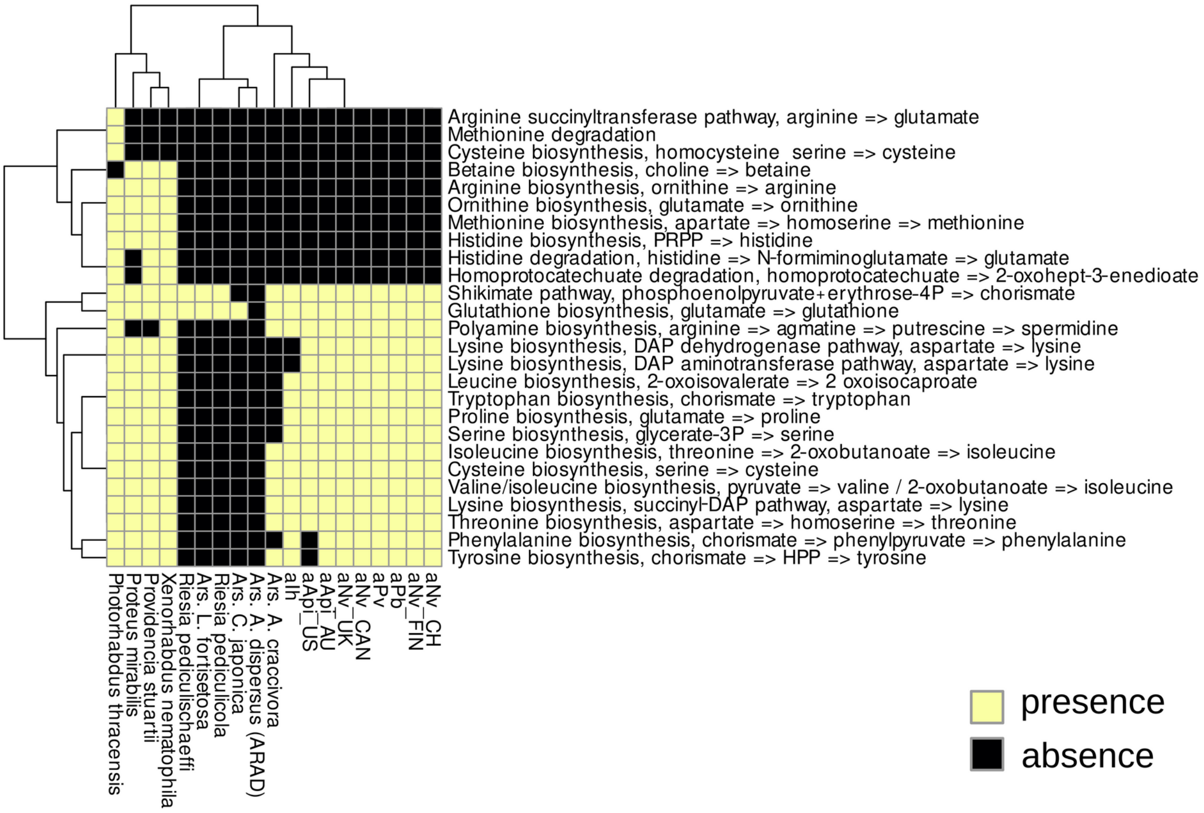


D) Metabolism of cofactors and vitamins

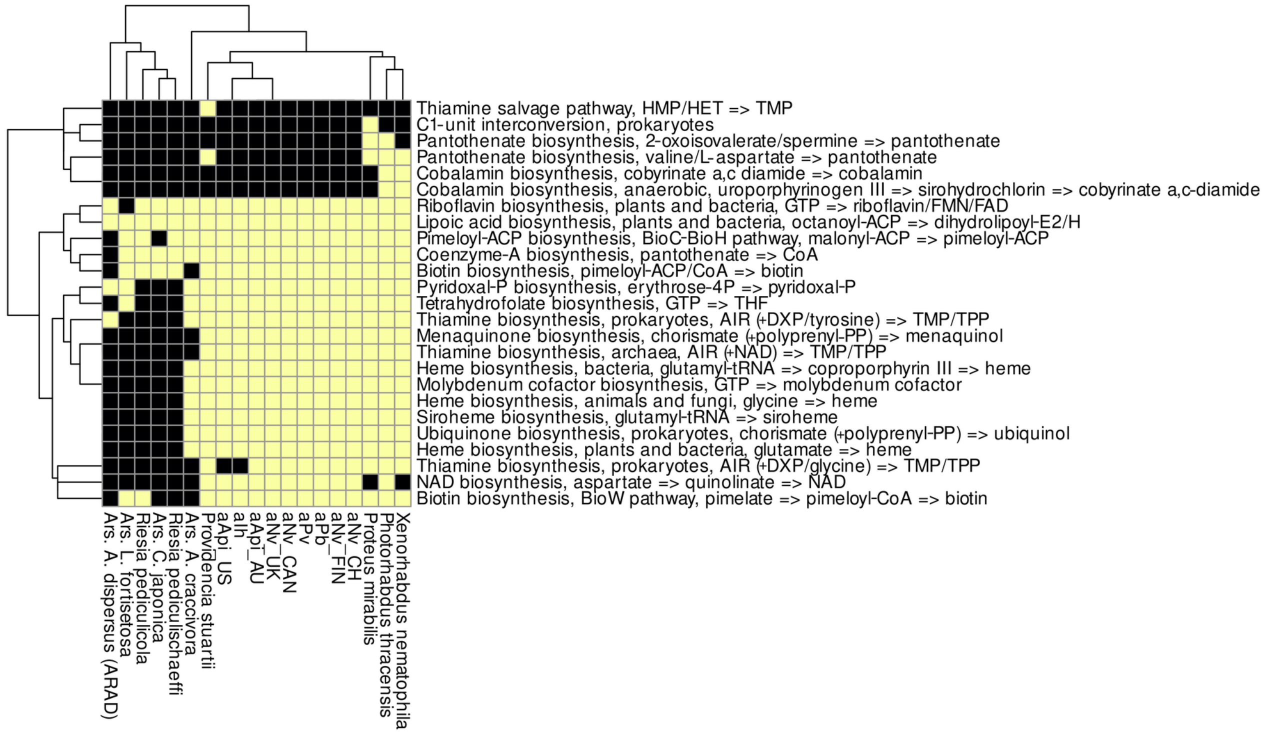

**Supplementary Figure S5** Comparison of **A**) the fraction of GC content and **B**) the average gene length (in amino acids) between core (common in all genomes) and non-core genes across the different *Arsenophonus* clades. Inset in (A) core section is a magnified version of the three main clades (apicola: purple, nasoniae: blue and triatominarum: yellow).

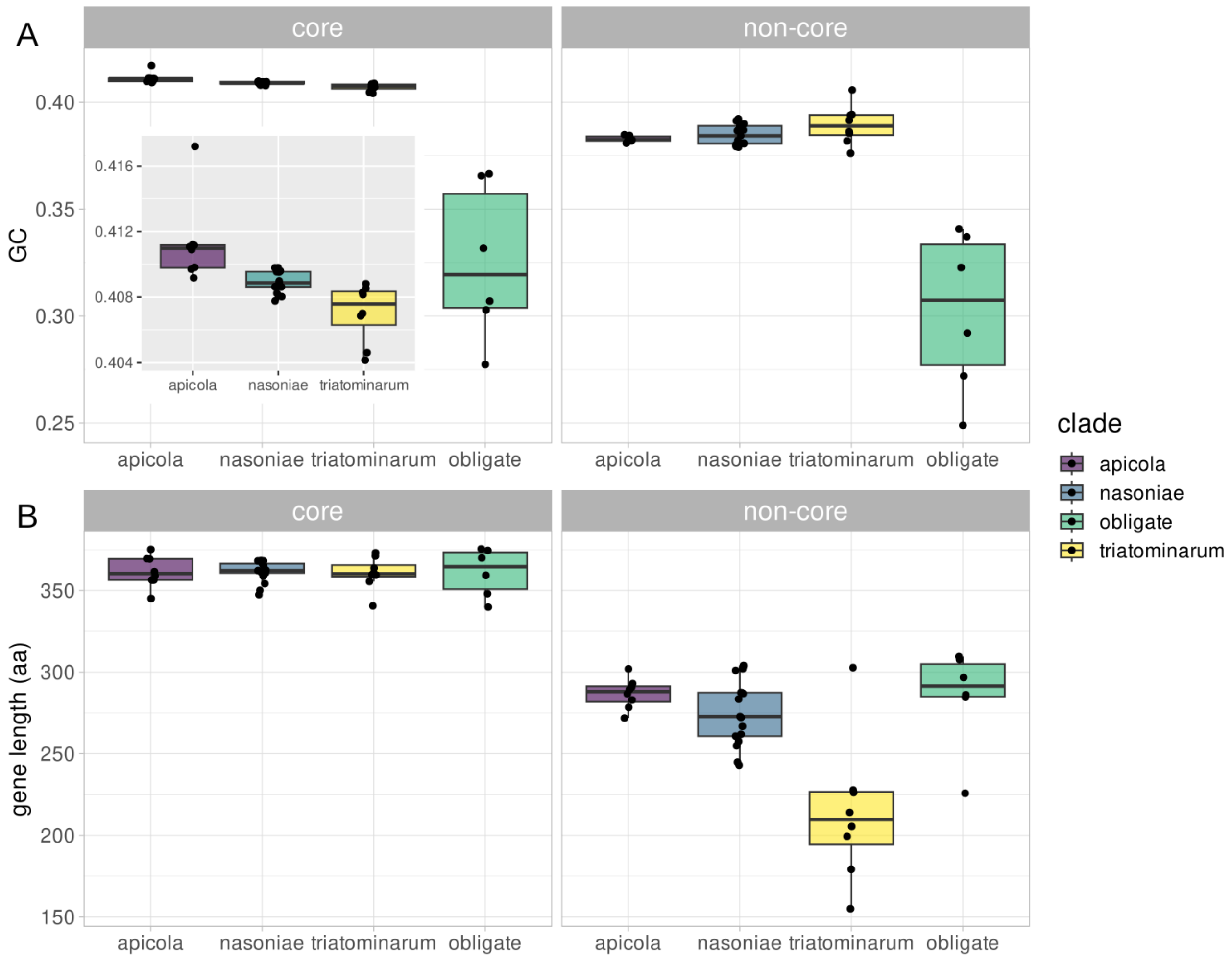

**Supplementary Figure S6** Association between genome size and the fraction of GC content for coding sequences (CDS) across the *Arsenophonus* genomes excluding the obligate strains. Circles: complete genomes, triangles: draft genomes. The red dashed line in panel B represents a fitted linear trend line with confidence intervals shown as grey shading.

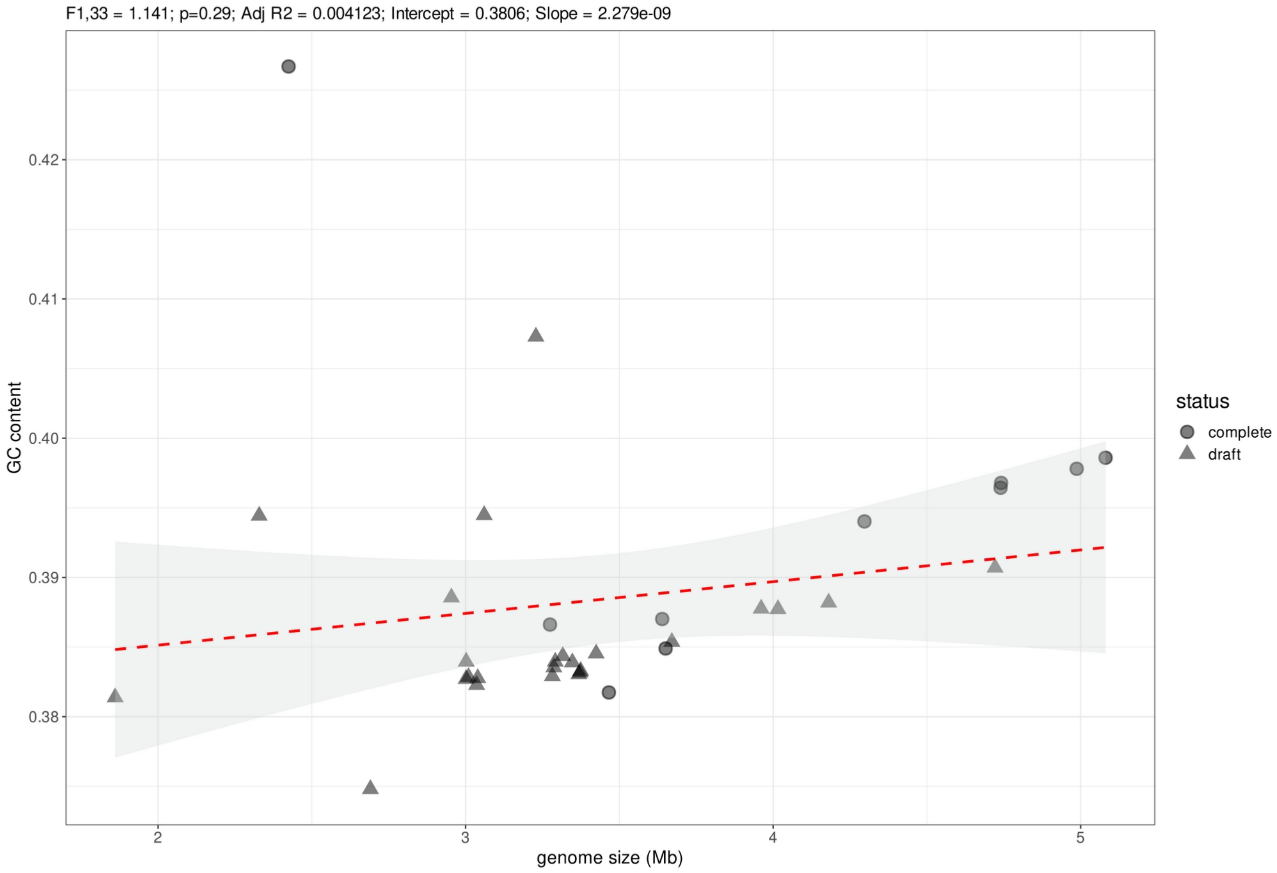

**Supplementary Figure S7** Predicted capacity at DNA repair genes and pathways across *Arsenophonus* genomes. Black: component absent, yellow: component present. Strain names in black: outgroup taxa, purple: apicola clade, blue: nasoniae clade, yellow: triatominarum clade and green: obligate strains.

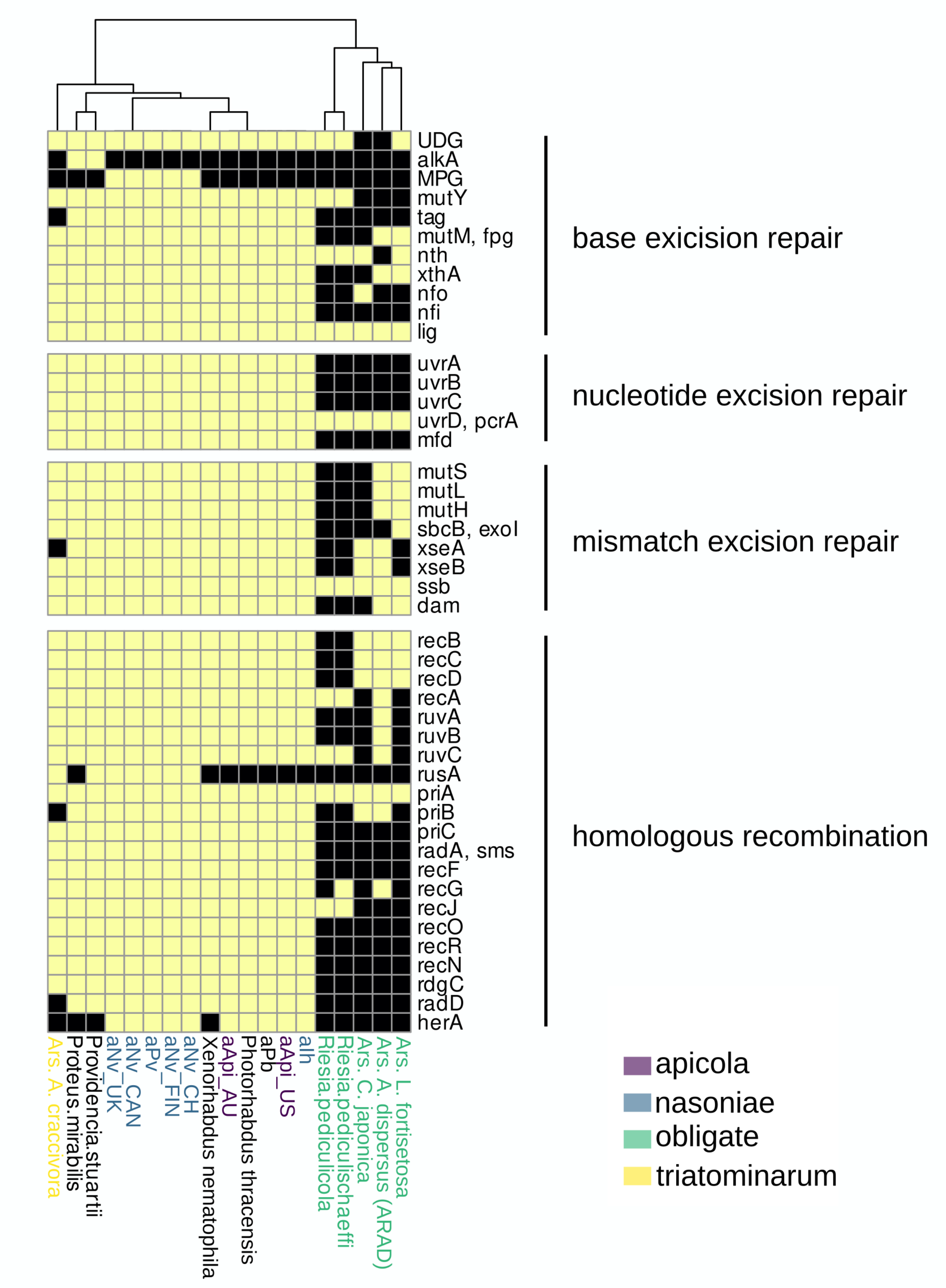

**Supplementary Figure S8** Phylogenetic cladogram showing the three sets of branches that were used for the analysis of relaxation of selection between Arsenophonus clades using RELAX in HyPhy Software.

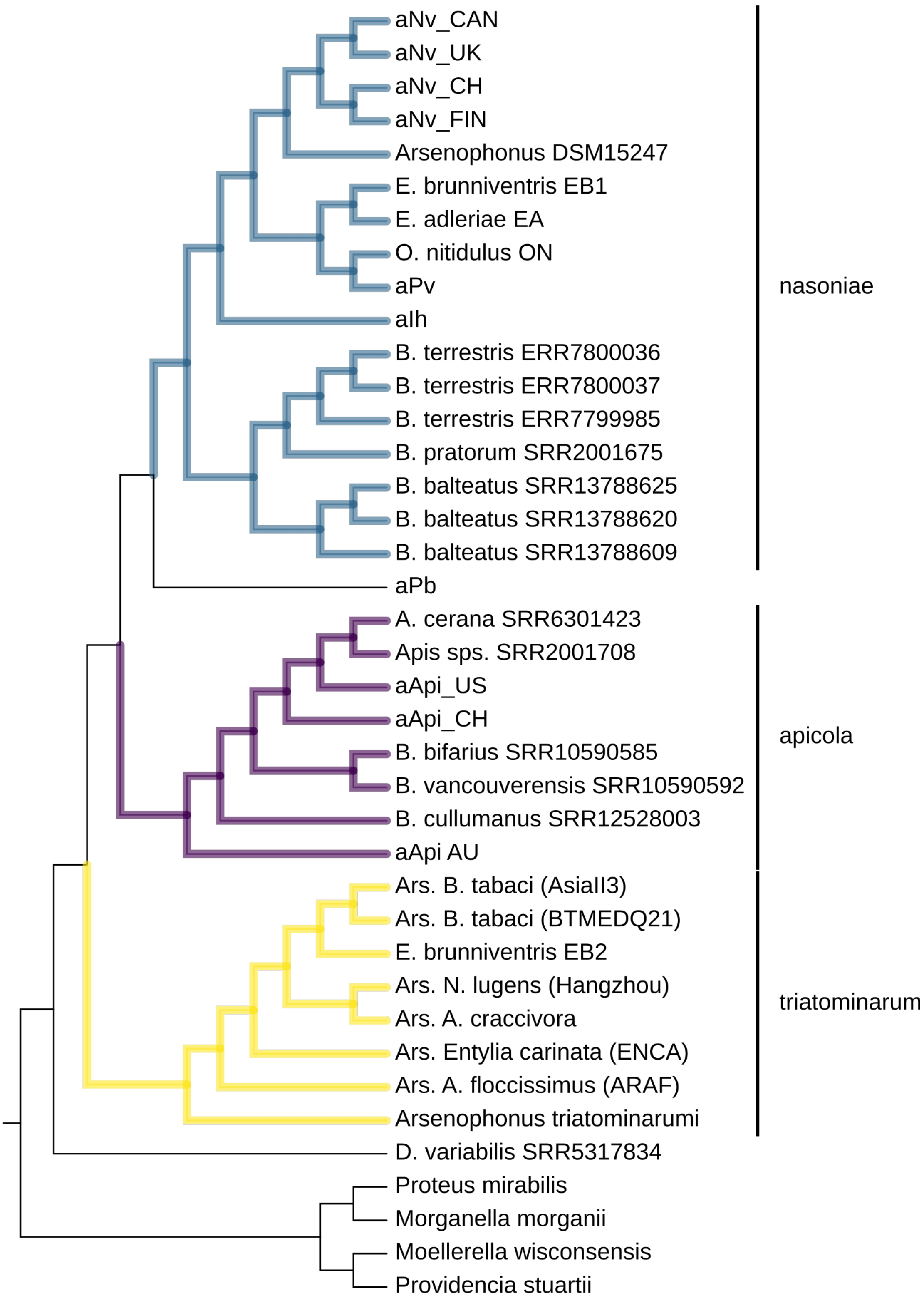
