## Supplementary Methods for "Genome dynamics across the evolutionary transition to endosymbiosis"

**Arsenophonus isolates and cultivation.**

*A. nasoniae* isolates aNv_UK, aNv_CH and aNv_CAN derived from *Nasonia vitripennis* from the UK, Switzerland and Canada respectively (isolation as previously described in [1, 2]. The isolation and culture of *A. nasoniae aPv* from *Pachycrepoideus vindemmiae* and the *Arsenophonus* strain *a*Pb from the butterfly *Polyommatus bellargus* is described in [3]. The *Arsenophonus nasoniae* strain ArsIxoH (*a*Ih) previously identified in the parasitoid wasp *Ixodiphagus hookeri* was isolated from questing *I. ricinus* nymphs collected in September 2020 by blanket dragging in De Buunderkamp, the Netherlands (52° 00′ N, 5° 44′ E). These nymphs were morphologically identified to species level using an identification key [4]. Ticks were kept live until processing. Isolation and culture of *A. nasoniae a*Ih was performed as described in [5] and harvested as described in [6] with modifications. The needle and syringe protocol was implemented using 26 gauge needles and a 0.45µm syringe-driven membrane filter. Isolation and culture of the *Arsenophonus apicola* strain *a*Api_AU from Australian honey bees is described in [7].

*Ca*. A. triatominarum was isolated from the host species *Triatoma infestans* maintained in the lab of Guenter Schaub (origin: colony 37, collected Bolivia 2005). The bacteria were isolated from surface sterilized *T. infestans* by collecting heamolymph. Two microliters of haemolymph was added to a 90% confluent cell monolayer of the *Drosophila melanogaster* S2 cell line (Drosophila Genetic Resource Centre stock number 6) grown in a 24-well plate with 1 ml of MMI plus 20% FBS. The preparation was centrifuged at 1,000x g at room temperature for 10 min to bring the bacteria into contact with the cells and incubated at 26.5°C for 16 h. At 10-day intervals, medium was taken from the cell layer and passaged onto a new 90% confluent S2 cell monolayer. The insect cell cultures were also tested routinely for microorganisms cultivable on 5% sheep blood agar plates incubated at 26.5°C, and they were inspected daily for 10 days at a magnification of x600 for microbial growth using an inverted M100 microscope (Swift-Microtec, Oxford, United Kingdom). Each symbiont-cell line was then cloned by limiting dilution, replicated three times and maintained as described above on S2 cell cultures. Cultures for genome sequencing were bulked up in 200 ml tissue flask. Total DNA was extracted from insects and 10-day-old insect cell cultures using a DNeasy tissue kit (QIAGEN, United Kingdom) following the manufacturer’s protocol for cultured animal cells.

**Sequencing, assembly, and annotation**

**Targeted sequencing and assembly of focal *Arsenophonus* strains**. All targeted genomes were sequenced using a combination of short (Illumina) and long (Nanopore) reads as described below, with the exception of *Ca. A. triatominarum*, which was completed solely with PacBio reads.

*Genome sequencing of Arsenophonus nasoniae strain a*Nv*_UK*. The *a*Nv_UK strain was sequenced following the same procedure as described in [8]. Briefly, high molecular weight (HMW) gDNA was prepared from a 50ml culture using a modified CTAB and phenol/chloroform extraction protocol [9]. Nanopore sequencing was performed with the Rapid Sequencing Kit (SQK-RAD004) (Oxford Nanopore, UK) on a FLO-MIN106 R9.4 MinION flow cell using 3ug of HMW gDNA and omitting the library loading beads to avoid blocking the sample port. The raw Nanopore reads were live basecalled in MinKNOW software v18.01.6 (Oxford Nanopore, UK). Low quality reads (quality score < 7) or small reads (<1kb) were discarded. Illumina sequencing was performed by MicrobesNG

(Birmingham, AL) using the Nextera XT library prep protocol on a MiSeq platform (Illumina, San Diego, CA, USA). Reads were adapter trimmed using Trimmomatic 0.30, with a sliding window quality cutoff of Q15 [10]. A hybrid assembly based on short and long reads was generated using the Unicycler pipeline version 0.4.5 under the normal mode [11]. The quality of the assembly was assessed by mapping the long reads back to it and manually inspecting in the Integrative Genomics Viewer (IGV) v2.8.9 for inconsistencies. A final round of polishing using the Illumina reads was performed with Polypolish v0.5.0 [12].

*Genome sequencing of the Arsenophonus nasoniae strains (aNv_CAN*, *aNv_CH* and *aPv*. These genomes were sequenced by MicrobesNG (Birmingham, UK) using their enhanced genome service. Briefly, long-read gDNA libraries were prepared with Oxford Nanopore SQK-RBK004 kit (Oxford Nanopore, UK) using 400–500 ng high molecular weight DNA and sequenced in a FLO-MIN106 (R.9.4.1) flow cell in a GridION (Oxford Nanopore, UK). Short-read Illumina sequencing was performed with the Nextera XT library prep protocol on a HiSeq platform (Illumina) using a 250 bp paired-end protocol. Reads were adapter trimmed using Trimmomatic 0.30, with a sliding window quality cutoff of Q15 [10]. An initial assembly of the long reads was performed using Flye assembler v2.8 under the uneven coverage mode ("-meta" option) and the "-plasmid" option enabled. Subsequently, the long reads were mapped back to each assembly and manually inspected for inconsistencies. Short read polishing of the assemblies was performed using five rounds of polishing with Pilon v1.22 [13] followed by a round of polishing with Polypolish v0.5.0 [12].

*Genome sequencing of the Arsenophonus strain aPb identified in the butterfly Polyommatus bellargus*. High molecular weight gDNA was extracted using a Qiagen genomic-tip 20/g and the Qiagen Genomic DNA protocol for Gram-negative bacteria (QIAGEN, UK) from about 1ml of liquid culture in Brain Heart Infusion (BHI) medium.  Long-read gDNA libraries were prepared with the Oxford Nanopore SQK-LSK109 kit (Oxford Nanopore, UK) using 500 ng high molecular weight DNA and sequenced on a FLO-FLGOP1 flow cell and the Flongle-MinION adapter (Oxford Nanopore, UK). Raw Nanopore reads were subsequently basecalled using Guppy v4.2.2 (Oxford Nanopore, UK) under the high accuracy model. A preliminary assembly was performed using Flye v2.8 under the uneven coverage mode ("-meta" option). Short Illumina reads obtained from Nadal-Jimenez et al. [3] were mapped to the long-read assembly using minimap2 v2.17-r941 [14] to identify and extract the *Arsenophonus* reads. Short and long *Arsenophonus* reads were used to prepare the final assembly using the Unicycler pipeline version 0.4.5 under the normal mode. Like previously, the quality of the final assembly was assessed by mapping the long reads back to it and manually inspecting for inconsistencies. A final round of polishing using the Illumina reads was performed with Polypolish v0.5.0 [12].

*Genome sequencing of Arsenophonus nasoniae strain aIh* Short Illumina reads were generated on the Illumina NovaSeq 6000 system (Illumina, San Diego, CA, USA) at Baseclear (Leiden, the Netherlands) from genomic DNA extracted using the ZymoBIOMICS™ 96 MagBead DNA Kit (Zymo Research, Orange, CA). High molecular weight gDNA was extracted using a Qiagen genomic-tip as described above for *Arsenophonus a*Pb. Long-read gDNA libraries were prepared with the Oxford Nanopore SQK-LSK109 kit (Oxford Nanopore, UK) using 1500 ng high molecular weight DNA and sequenced in a FLO-MIN106D (R.9.4.1) MinION flow cell (Oxford Nanopore, UK). Raw Nanopore reads were subsequently basecalled using Guppy v4.2.2 (Oxford Nanopore, UK) under the high accuracy model. Low quality and short reads were removed and the remaining reads were assembled using the Flye assembler v2.8 under the uneven coverage mode ("-meta" option) and the "-plasmid" option enabled. The quality of the assembly was assessed as aforementioned by mapping the long reads back to the assembly and manually inspected for inconsistencies. Short read polishing was performed using five rounds of polishing with Pilon v1.22 [13] followed by a round of polishing with Polypolish v0.5.0 [12].

*Genome sequencing of Arsenophonus apicola strain aApi_AU*. This genome was sequenced by Charles River Laboratories (Australia). In brief, high molecular weight DNA was extracted and Long-read gDNA libraries were prepared with the Oxford Nanopore SQK-LSK110 kit (Oxford Nanopore, UK) which were then sequenced on a single FLO-MIN106D (R.9.4.1) MinION flow cell (Oxford Nanopore, UK). Long reads >1kb were used for genome assembly using the Flye assembler v2.9.1 under the uneven coverage mode ("-meta" option). Assembly QC and short read polishing was performed as described above; Illumina reads were derived from the previous study [7].

*Genome sequencing of Ca. A. triatominarum*: The high quality draft genome of "Ca. Arsenophonus triatominarum" was generated using PACBIO long reads from four SMRT cells (C2, P4 chemistry). The reads were filtered using BLAST to remove *Drosophila* reads and assembled with HGAP using default parameters [15]. The assembly yielded 94 contigs corresponding to the *Arsenophonus* genome, spanning the total length of 3,858,720 bp with 70x fold average coverage.
